# BamToCov: an efficient toolkit for sequence coverage calculations

**DOI:** 10.1101/2021.11.12.466787

**Authors:** Giovanni Birolo, Andrea Telatin

## Abstract

Many genomics applications requires the calculation of nucleotide coverage of a reference or counting how many reads maps in a reference region.

Here we present BamToCov, a suite of tools for rapid and flexible coverage calculations relying on a memory efficient algorithm and designed for flexible integration in bespoke pipelines. The tools of the suite will process sorted BAM or CRAM files, allowing to extract coverage information using different filtering approaches.

BamToCov tools, unlike existing tools already available, have been developed to require a minimum amount of memory, to be easily integrated in workflows, and to allow for strand-specific coverage analyses. The unique coverage calculation algorithm makes it the ideal choice for the analysis of long reads alignments. The programs and their documentation are freely available at https://github.com/telatin/bamtocov.

## 1 Introduction

Sequencing coverage calculations have been done since the dawn of genomics (Lander and Waterman, 1988), commonly in relation to *a priori* theoretical calculations aimed at understanding the amount of effort required to produce sufficient DNA reads with capillary sequencers.

With the advent of massively parallel sequencing (also referred to as ‘next generation sequencing’) those *a priori* calculations began to be matched by *a posteriori* calculations made by mapping the DNA reads against a reference sequence (either a pre-existing reference, or the *de novo* assembly of the sequencing output itself). In this context some bases sequenced would not be accounted for (*e. g*. adaptor sequences, contaminants, or unmappable reads).

When using paired libraries, it is also possible to evaluate the *physical coverage, i. e*. the number of times a base is spanned by a read pair.

There are already several tools to extract coverage information from alignment files (in BAM format): Samtools (Li *et al*., 2009), Bedtools (Quinlan, 2014), Sambamba (Tarasov *et al*., 2015) and the newer and more feature-rich Mosdepth (Pedersen and Quinlan, 2018b) and MegaDepth (Wilks *et al*., 2021). A common limitation of the existing tools is the inability to calculate physical coverage which is important when determining the integrity of assemblies using mate-pairs libraries. Also, it is not possible to separate the coverage per strand; if a position is covered only by forward reads, or only by reverse reads, it is probably due to misalignment. To address these limitations, we developed Covtobed (Birolo and Telatin, 2020), a simple yet efficient C++ program which, inspired by the UNIX philosophy of computer programming, focused on a single task supporting input and output streams. Here we introduce BamToCov program and its auxiliary utilities, written in the Nim language; this performs coverage calculations using the core algorithm of Covtobed with new features to support interval targets, new output formats, coverage statistics and multiple BAM files, while retaining the ability to read input streams, thereby achieving an overall performance improvement (*i. e*. a smaller memory footprint and an increase in speed of up to 3×).

### Sequence coverage and physical coverage

When using paired libraries, we can also evaluate the physical coverage, the average number of times a base is spanned by a read pair. Figure 1 depicts the difference between sequence and physical coverage.

**Figure 1.**
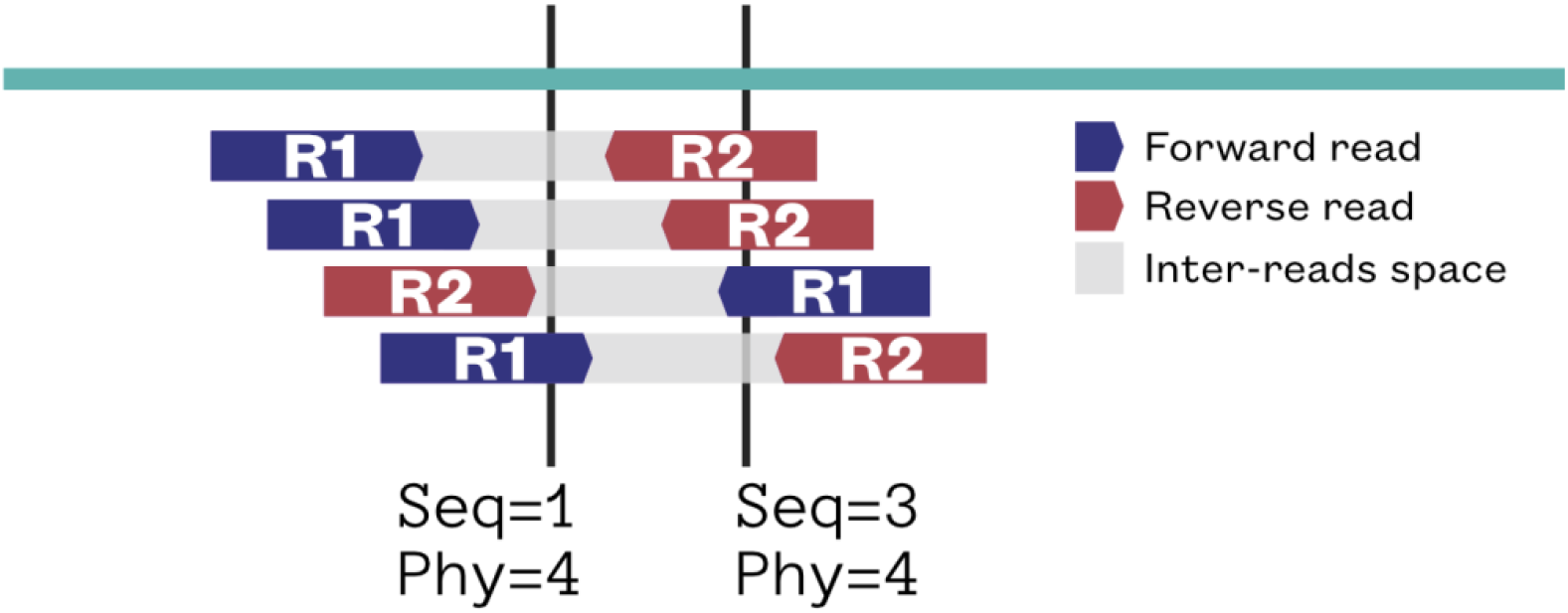
Schematic representation of paired-end reads (blue and purple arrows) mapped against a reference sequence (red). The sequence coverage (Seq) and the physical coverage (Phy) of two nucleotide coordinates (black lines) have been calculated, showing the difference between the two.

The analysis of physical coverage allows to understand if the library, rather than the sequencing reads, sampled a region. If the physical coverage drops unexpectedly, this can be an indication of a misassembled region, hence the possibility to quickly identify variations in its value can be beneficial.

If the sequencing library preparation is supposed to be strand unbiased, then a strand bias in the alignment can be an indicator of a mapping problem.

## 2 Methods

BamToCov implements the coverage calculation algorithm of Covtobed, a C++ program, but using hts-lib (Bonfield et al., 2021) for BAM/CRAM parsing, via the hts-nim wrapper (Pedersen and Quinlan, 2018a) instead of libbamtools, hence natively supporting CRAM files.

The programs have been written in Nim and tested using three compiler versions (1.0, 1.2, and 1.4). Rather than using a vector of integers having the size of the sequence, BamToCov uses a streaming approach that requires sorted alignments as input, computing coverage is computed starting from zero at the leftmost base in each contig and updated on-the-fly while reading alignments and moving toward the right. Coverage is incremented at the start of each alignment and decremented at the end, keeping the ending positions in a priority queue.

The suite of tools is automatically tested, and available via the BioConda project (The Bioconda Team et al., 2018).

The scripts used to benchmark the execution times and the peak memory usage are available in the software repository.

## 3 Results

BamToCov can be used as a drop-in replacement for Covtobed, and we carefully tested the consistency of the results and benchmarked the speed improvement of the new implementation if compared with the original.

We also compared the performance of BamToCov with the available packages performing the same calculation, highlighting that BamToCov is second only to MegaDepth in terms of speed.

### Features

Building upon the design of Covtobed, BamToCov is designed to support input streams (hence does not require indexed BAM files), and to produce physical coverage and per-strand coverage calculations.

BamToCov also allows the user to provide a set of intervals of interest (target), used both to produce an output BED file restricted to those regions, and generate a table of statistics per each interval.

Supplementary utilities allow to focus on read counts, rather than sequence coverage, or statistics over the whole chromosomes.

BamToCov supports targets in BED, GFF and GTF formats.

### Performance

To evaluate the performance of BamToCov, we adopted four test datasets with different size, density of alignments and different read length:

1. Fungus, SR: Alignment of a whole genome shotgun of Short Reads (SR), produced via Illumina MiSeq sequencing, against the assembly of an isolate of Candida albicans
2. Fungus, LR: Alignment of a whole genome shotgun of Long Reads (SR), produced via ONT Nanopore sequencing, against the assembly of an isolate of Candida albicans
3. Exome: Illumina sequencing of sample HG00258 from the 1000 Genomes project (Homo sapiens)
4. Gene Panel: Illumina sequencing of a target enrichment using a panel of 16 genes (Homo sapiens)

BamToCov is a fast program, up to 2X faster than its predecessor (CovToBed), and very promising when evaluating the coverage of long reads (see Table 1).

**Table 1.**
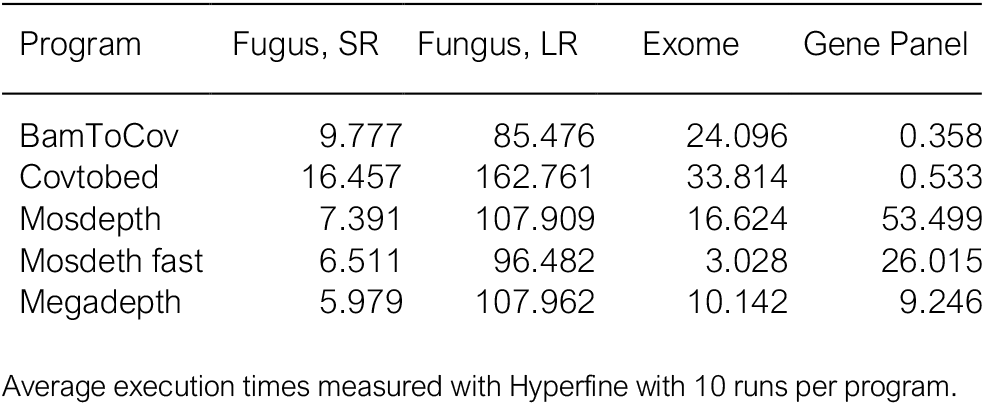
Execution time in seconds

BamToCov is the most memory efficient program for coverage calculations, with an improvement up to 50% if compared with the C++ implementation (Covtobed). Other programs relying on chromosome-sized vectors require up to 400X more memory to analyse a typical human exome (Table 2). BamToCov memory usage is not affected by the genome size.

**Table 2.** Peak memory usage

| Program | Fugus, SR | Fungus, LR | Exome | Gene Panel |
| --- | --- | --- | --- | --- |
| BamToCov | 2,740 | 4,376 | 5,700 | 2,172 |
| Covtobed | 4,080 | 5,008 | 6,588 | 4,052 |
| Mosdepth | 13,952 | 19,140 | 1,983,928 | 6,425,744 |
| Megadepth | 11,644 | 11,636 | 995,232 | 980,040 |

Peak memory usage in bytes

BamToCov provides the most memory efficient algorithm for coverage calculation, that consistently outperformed alternative programs in four test datasets (Table 2).

## 4 Conclusion

BamToCov is a program and suite of utilities engineered to simplify their application in bioinformatics pipelines requiring coverage calculations. It is designed for the needs of bioinformaticians working in microbial genomics and non-model organisms, where the possibility of working from streams can be particularly beneficial and is unsupported by other comparable tools.

The peculiar algorithm adopted is the most memory efficient by far for larger genomes, and the new implementation in Nim yields further performance benefits both in terms of execution times and memory footprint.

## Acknowledgements

The authors express their gratitude to Rebecca Ansorge and Stefano Romano, for testing the suite.

